# Proteomic analysis reveals the dominant effect of ipomoeassin F on the synthesis of membrane and secretory proteins in triple-negative breast cancer cells

**DOI:** 10.1101/2024.07.28.605505

**Authors:** Brihget Sicairos, Jianhong Zhou, Zhijian Hu, Qingyang Zhang, Wei Q Shi, Yuchun Du

**Affiliations:** Department of Biological Sciences, University of Arkansas, Fayetteville, Arkansas 72701, USA; Feinstein Institute for Medical Research, Northwell Health, 350 Community Dr., Manhasset, New York, 11030, USA; Department of Mathematical Sciences, University of Arkansas, Fayetteville, Arkansas 72701, USA; Department of Chemistry, Ball State University, Muncie, Indiana 47306, USA

## Abstract

Ipomoeassin F (Ipom-F) is a natural compound with embedded carbohydrates that exhibits a potent cytotoxic effect on triple-negative breast cancer (TNBC) cells. The mechanism behind this selective potency remains unclear. To elucidate this mechanism, we analyzed the proteome profiles of the TNBC MDA-MB-231 cells after exposure to Ipom-F at different time points and increasing doses using a quantitative proteomic method. Our proteomic data demonstrate that the major effect of Ipom-F on MDA-MB-231 cells is the inhibition of membrane and secreted protein expression. Our proteomic data are consistent with the recently uncovered molecular mechanism of action of Ipom-F, which binds to Sec61-α and inhibits the co-translational import of proteins into the endoplasmic reticulum. We have defined a subset of membrane and secreted proteins particularly sensitive to Ipom-F. Analysis of the expression of these Ipom-F-sensitive proteins in cancer cell lines and breast cancer tissues demonstrates that some of these proteins are upregulated in TNBC cells. Thus, it is likely that TNBC cells may have adapted to the elevated levels of some proteins identified as sensitive to Ipom-F in this study; inhibition of the expression of these proteins leads to a crisis in proliferation and/or survival for the cells.

## Introduction

Natural products are a major resource for drug development and translational biomedical research. Ipomoeassins are a family of plant-derived macrolides with embedded carbohydrates. Among the six members, ipomoeassin F (Ipom-F) demonstrated the most potent cytotoxic effect on several cancer cell lines with single-digit nanomolar IC50s (1–3). The natural abundance of ipomoeassins is generally low, except for the less active ipomoeassin A. This scarcity has historically limited research on this promising class of natural products (2). After successful total synthesis of Ipom-F (4, 5) and particularly after we identified Sec61-α as the molecular target of Ipom-F in human cells (6), rapid progress has been made. Subsequent studies, both *in vitro* and *in vivo*, have confirmed that Ipom-F plays a key role in regulating the synthesis of membrane-related proteins by inhibiting the co-translational import of those proteins into the endoplasmic reticulum (ER) membrane or ER lumen during protein synthesis (7–11). Recently, the structure basis for the inhibitory effect of Ipom-F on Sec61-α has been established (12).

Ipom-F is a highly potent cytotoxic natural product with IC50s of low nM for many cancer cell lines (13). Although the molecular target of Ipom-F, Sec61-α, is ubiquitously required for synthesizing membrane and secretory proteins in most eukaryotic cells, our previous studies demonstrated that Ipom-F differentially affected the viability of different cancer cells. Specifically, when Ipom-F was tested on the NCI-60 human tumor cell lines, Ipom-F exhibited potent cytotoxicity toward triple-negative MDA-MB-231 breast cancer cells and a few other cancer cell lines, while having low or moderate cytotoxicity toward some other cancer cell lines (7, 13). The mechanism behind the high cytotoxic potency of Ipom-F toward the triple-negative breast cancer (TNBC) cells remains unclear. To address this issue, we analyzed the proteome profiles of TNBC MDA-MB-231 cells after treating the cells with Ipom-F at different time points (time course) and increasing doses (dose curve) using a quantitative proteomic method. Consistent with the molecular mechanism of action of Ipom-F in cells established in our biochemical studies (6), our proteomic data demonstrated that the major effect of Ipom-F on MDA-MB-231 cells was to inhibit the expression of membrane and secretory proteins. We have identified a subset of membrane and secretory proteins that were particularly sensitive to Ipom-F. When we compare the protein levels of some of the Ipom-F-sensitive membrane and secreted proteins among different cancer cell lines and breast cancer tissues, it was evident that the levels of some Ipom-F-sensitive proteins were significantly higher in MDA-MB-231 cells compared to others. Thus, it is likely that the high cytotoxicity of Ipom-F toward TNBC MDA-MB-231 cells is due to the reliance of the cells on the elevated levels of some of the membrane and secretory proteins, which were identified to be particularly sensitive to Ipom-F in this study. Interestingly, our proteomic analysis revealed that the levels of MHC class I and MHC class II proteins were significantly inhibited by Ipom-F. This suggests that Ipom-F may act as an immunosuppressive agent, a feature indicated in our previous study (6) but not yet well explored.

## Materials and Methods

### Cell culture, proteome labeling, and Ipom-F treatments

MDA-MB-231 cells were routinely maintained in Dulbecco’s modified Eagle’s medium (DMEM) supplemented with 10% fetal bovine serum (FBS), and 1% penicillin and streptomycin (Invitrogen, CA). We used a SILAC (stable isotope labeling by amino acids in cell culture)-based quantitative proteomic method (14, 15) to identify and quantify the proteins that were differentially expressed in the MDA-MB-231 cells after the cells were treated for different periods of time at a fixed dose of Ipom-F (time course) or with different doses of Ipom-F for a fixed period of time (dose curve). SILAC proteome labeling was conducted as described previously (7, 16–18). Briefly, the proteome of the MDA-MB-231 cells was isotopically labeled for two weeks by growing the cells in DMEM containing either unlabeled arginine and lysine (“light”), ^13^C_6_-arginine and ^2^H_4_-lysine (“medium”), or ^13^C_6_ ^15^N_4_-arginine and ^13^C_6_ ^15^N_2_-lysine (“heavy”) supplemented with 10% dialyzed FBS. The isotopically labeled cells were then used for the Ipom-F time course and dose curve studies. For the time course, one set of the “light,” “medium,” and “heavy” cells was treated with 18 nM Ipom-F for 0, 4 or 8 hours, and a second set of the “light”, “medium”, and “heavy” cells was treated with 18 nM Ipom-F for 0, 12 or 16 hours. For the dose curve, one set of the “light,” “medium,” and “heavy” cells was treated with 0, 3 or 6 nM Ipom-F for 11 hours, and a second set of the “light”, “medium”, and “heavy” cells was treated with 0, 18 or 54 nM Ipom-F for the same period. After the treatments, the cells were lysed, and the total protein was prepared for LC-MS/MS as described previously (7, 16–18).

### SDS-PAGE, LC-MS/MS, and data analysis

In-gel digestion, database search, and LC-MS/MS data quantification were performed as described previously (7, 17, 19). Specifically, equal amounts of the “light,” “medium,” and “heavy” total protein from each set of the time course or dose curve experiments (40 µg/each) were mixed. The mixed proteins (120 µg) were then fractionated by a 12% SDS-PAGE (15.5 cm x 18 cm), followed by Coomassie Brilliant Blue staining. Each protein lane (the entire lane) of the Coomassie brilliant Blue-stained gel was cut into 12 slices of equal size, and the gel slices were subjected to in-gel digestion. The resulting peptides were analyzed by LC-MS/MS using an Orbitrap Fusion mass spectrometer (ThermoFisher, San Jose, CA) operated in a data-dependent mode for tandem MS as described previously (17, 20, 21).

Raw data from the LC-MS/MS analysis were processed by MaxQuant (version 1.6.1.0) (22, 23) with the built-in search engine Andromeda (24) and searched against a target-decoy (25) human SwissProt protein database (October, 2023) retrieved from UniProt (www.uniprot.org) (26). The false discovery rates (FDRs) for peptide and protein identification were set to 1%. The MS error tolerance was set to 4.5 ppm, and the MS/MS error tolerance was set to 0.5 Da. The minimum required peptide length was set to 7 amino acids, and a maximum of 2 missed cleavages was allowed. The variable modifications of ^13^C_6_-arginine, ^2^H_4_-lysine, ^13^C_6_ ^15^N_4_-arginine, ^13^C ^15^N -lysine, oxidation of methionine and protein N-terminal acetylation, and the fixed modification of cysteine carbamidomethylation were included. SILAC ratios (heavy/light and medium/light ratios) were calculated using unique and razor peptides with a minimum ratio count of 2 (23). The proteins matched to the reverse database, identified only by site or single peptide, and the common contaminants were discarded. Protein expression changes at different time points (4, 8, 12, or 16 hours after treatment) and at increasing Ipom-F concentrations (3, 6, 18, or 54 nM) were calculated by comparing the peptide signal intensities in LC-MS/MS between each time point or Ipom-F concentration and their respective controls (0 hours and 0 nM treatment). Proteins with SILAC ratios ≤ 0.8 or ≥ 1.2 at each time point were selected, and the selected proteins were compared across the time course to identify the proteins consistently affected by Ipom-F. Proteins consistently downregulated by at least 20% (SILAC ratios ≤ 0.8) or upregulated by at least 20% (SILAC ratios ≥ 1.2) after Ipom-F treatments across the four time points were defined as Ipom-F-regulated proteins. The same approach was used to identify the Ipom-F-regulated proteins in the studies on increasing Ipom-F concentrations.

### Bioinformatics analysis of LC-MS/MS data

Enrichment analyses of cellular components, biological processes, and molecular functions were performed on the Ipom-F-regulated proteins using FunRich, a stand-alone software tool for functional enrichment and interaction network analysis of genes and proteins (27, 28) or the Database for Annotation, Visualization, and Integrated Discovery (DAVID), a comprehensive set of functional annotation tools for investigators to understand the biological meaning behind large lists of genes (29, 30). Only the Ipom-F-inhibited proteins were analyzed for enrichment of cellular components, biological processes, and molecular functions because the number of Ipom-F-upregulated proteins was not sufficient enough for these analyses. In the FunRich analysis, the cellular components and molecular functions with p-values ≤ 0.05 were significantly enriched in the Ipom-F-inhibited proteins. In DAVID, we used the Ipom-F-inhibited proteins to probe the GOTERM_BP_DIRECT and GOTERM_MF_DIRECT databases to identify the enriched biological processes and molecular functions. Annotation clusters with enriched terms with p-values ≤ 0.05 were considered significantly enriched.

### Identification of proteins that are particularly sensitive to Ipom-F

To identify the proteins particularly sensitive to Ipom-F, we focused on the proteins whose expression was inhibited at least 25% by Ipom-F shortly after Ipom-F treatment (i.e., 4 and 8 h) and at lower Ipom-F concentrations (i.e., 3 and 6 nM). An overlap analysis of the Ipom-F-sensitive proteins identified by the time course experiments and the dose course experiments was carried out to identify the proteins consistently regulated by Ipom-F across the four separate studies, and the overlapping proteins were defined as Ipom-F-sensitive proteins.

The proteomics data for each of the identified Ipom-F-sensitive proteins across the NCI-60 human tumor cell lines were downloaded from the Cancer Dependency Map (DepMap) portal (https://depmap.org/portal/) (31). The downloaded proteomics data was analyzed using the heatmapper web tool (http://heatmapper.ca/expression/) to identify clusters of cell lines based on the expression levels of the Ipom-F-sensitive proteins (32). Briefly, we used average linkage for the hierarchical clustering and Spearman’s rank correlation for the clustering distance in the clustering analysis (32). In this context, average linkage calculates the average of all the distances between any two points to determine suitable clustering in hierarchical clustering. Spearman’s rank correlation measures the strength and the direction of the association between two ranked variables.

We also examined the levels of several Ipom-F-sensitive proteins available on the University of Alabama at Birmingham Cancer data analysis website (<u>UALCAN (uab.edu)</u>) (33, 34) in different subtypes of breast cancer and normal tissues. The UALCAN website provides protein expression analysis options using data from the Clinical Proteomic Tumor Analysis Consortium (CPTAC) and the International Cancer Proteogenome Consortium (ICPC) datasets (33). In the UALCAN website, the protein levels are presented in Z-values which represent the standard deviations from the median across samples. The CPTAC Log2 spectral count ratio values were normalized within each sample and across samples. The differential expression was considered significant if the p-value was ≤ 0.05.

### Western blotting

MCF-7 and MDA-MB-231 cells were treated with 1 or 5 nM Ipom-F for 12 hours. The cells were lysed, and the proteins were separated on a 12% SDS-PAGE (10 cm x 8 cm) at 170 V for 50 minutes and then transferred onto a cellulose membrane at 260 mA for 70 minutes. The remaining Western blotting procedures were conducted as described previously (18, 35). The anti-CD74, anti-CST1, and anti-CCN1 antibodies were purchased from Proteintech (Rosemont, IL). The anti-tubulin antibodies were purchased from Santa Cruz Biotechnology (Dallas, TX).

## Results and Discussion

### Identification of the Ipom-F-regulated proteins

To identify the proteins affected by Ipom-F in MDA-MB-231 cells, we cultured the cells in unlabeled and SILAC-labeled media. We then mock-treated the cells or treated them with 18 nM Ipom-F for 4, 8, 12, and 16 hours to perform a time course study. Additionally, we mock-treated or treated the cells with 3, 6, 18, and 54 nM Ipom-F for 11 hours to perform a dose curve study. Protein expression at each time point or dose was represented by the SILAC ratios of the Ipom-F treated cells (4, 8, 12, and 16 hours or 3, 6, 18, and 54 nM) relative to their respective controls (0 hours or 0 nM) (Supplementary Tables 1 and 2). The proteins with SILAC ratios ≤ 0.8 or ≥ 1.2 at each time point were selected. The selected proteins at each time point were then examined across the time course to determine the proteins that were consistently affected by Ipom-F. The proteins that were consistently downregulated by at least 20% (SILAC ratios ≤ 0.8) or upregulated by at least 20% (SILAC ratios ≥ 1.2) in response to Ipom-F across the four time points were defined as Ipom-F-regulated proteins. We performed a similar analysis for the data on the dose curve study. The time course analysis identified 84 proteins with lower expression compared to only 14 proteins with higher expression consistently across the time course (Supplementary Table 3). In comparison, the dose curve analysis identified 54 proteins with lower expression compared to only 11 proteins with higher expression consistently across the increasing Ipom-F concentrations (Supplementary Table 4). These results demonstrated that the predominant effect of Ipom-F in MDA-MB-231 cells was to inhibit rather than enhance protein expression.

To understand the types of proteins regulated by Ipom-F, we analyzed the Ipom-F-regulated proteins (Supplementary Tables 3 and 4) with FunRich (27, 28) and DAVID (29, 30). The enrichment analyses of the Ipom-F-downregulated proteins identified in the time course analysis revealed that proteins associated with cellular and organelle membranes, organelles (e.g., lysosomes, mitochondria, and Golgi), and secreted and vesicular proteins (e.g., extracellular proteins and exosomes) were significantly enriched (Fig. 1A and Supplementary Table 5). Enrichment analyses of the Ipom-F-downregulated proteins identified in the dose curve study showed similar results (Fig. 1B and Supplementary Table 6). The number of Ipom-F-upregulated proteins in the time course or dose curve studies was not sufficient enough for enrichment analyses with FunRich or DAVID. Our previous studies have shown that Ipom-F binds Sec6-α (6) and inhibits Sec61-α-mediated co-translational import of membrane proteins and secreted proteins into the endoplasmic reticulum (ER) (7–9). The proteomic results in this study are consistent with the molecular mechanism of action of Ipom-F reported in the previous studies.

**Figure 1.**
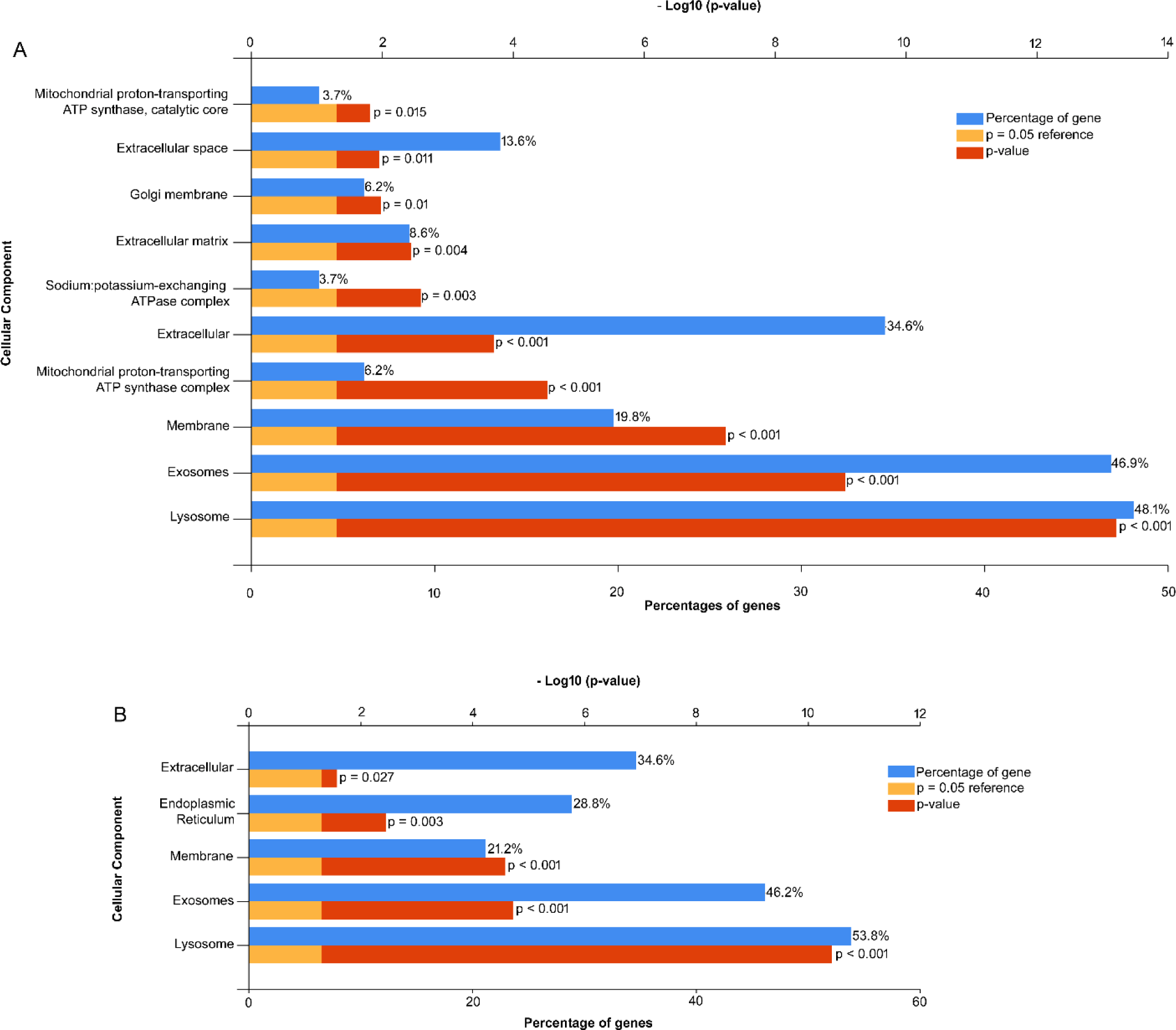
Ipom-F-inhibited proteins are membrane/organelle associated proteins or secreted proteins. The proteins identified to be consistently downregulated by Ipom-F across time points (**A**) and increasing doses of Ipom-F (**B**) were analyzed for enrichment of cellular components using FunRich. Only the components with a p-value ≤ 0.05 are displayed.

### Identification of the proteins that are particularly sensitive to Ipom-F

Since MDA-MB-231 cells are very sensitive to Ipom-F treatment (7, 13), Ipom-F likely targets the Achilles’ heel of the cells. We reasoned that the proteins responsible for the vulnerability of MDA-MB-231 cells to Ipom-F must be particularly sensitive to Ipom-F; therefore their expression must be affected by Ipom-F at earlier time points or lower doses of Ipom-F. Thus, we focused on the proteins that were affected by Ipom-F at earlier time points (4 and 8 hours) in the time course studies and at lower doses (3 nM and 6 nM) in the dose curve studies. For these analyses, we used a SILAC ratio ≤ 0.75 to identify the proteins that were particularly sensitive to Ipom-F; this parameter was slightly more stringent than the SILAC ratio used for the general identification of the proteins whose expression was affected by Ipom-F (Supplementary Tables 3 and 4). In the time course studies, the expression of 140 and 114 proteins was inhibited by at least 25% after 4 and 8 hours of Ipom-F treatment compared to the control (e.g., 0 h), respectively. When these two lists of proteins were compared, 78 proteins were identified in both lists, representing those particularly sensitive to Ipom-F in the time course analysis. In the dose curve studies, the expression of 88 and 119 proteins was inhibited by at least 25% following 3 and 6 nM Ipom-F treatments for 11 hours compared to their control (e.g., 0 nM), respectively. When these two lists of proteins were compared, 52 proteins were identified in both lists, representing those particularly sensitive to Ipom-F in the dose curve analysis. Overlapping analysis of the 78 proteins identified in the time course studies and the 52 proteins identified in the dose curve studies resulted in 12 proteins found in both lists (Fig. 2A and Table 1). We defined these 12 proteins as Ipom-F-sensitive proteins with high confidence because Ipom-F inhibited the expression of these proteins at earlier time points and lower doses, and this inhibition was consistently observed in four independent studies. According to the UniProt database (26), all 12 proteins were secretory proteins or membrane/organelle-related proteins (Table 1).

**Figure 2.**
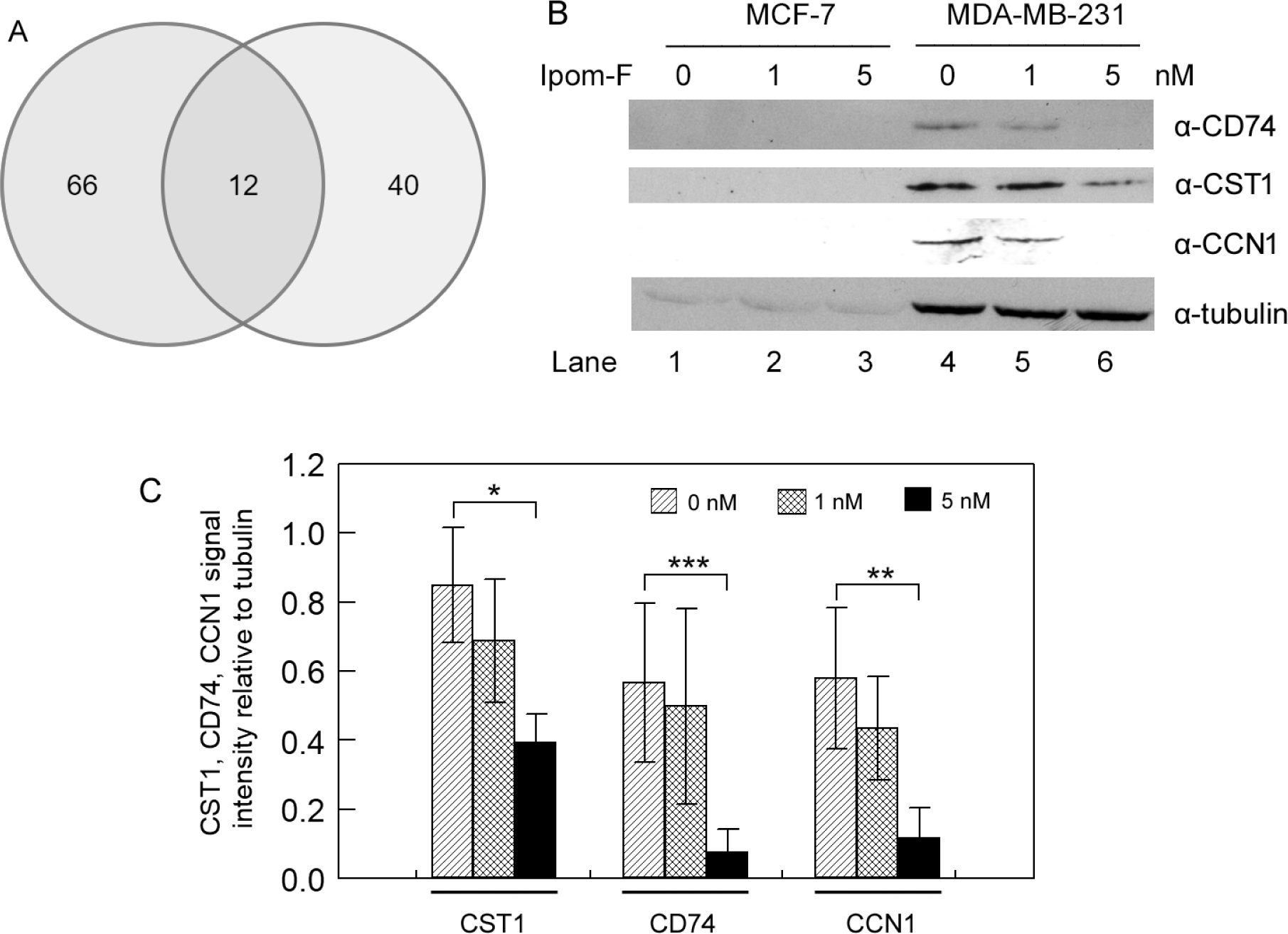
Identification and validation of proteins particularly sensitive to Ipom-F. (**A**) Overlap analysis of Ipom-F-sensitive proteins identified in the time-course and dose-curve studies. (**B**) Western blot analysis of the Ipom-F-sensitive proteins CD74, CST1, and CCN1 in MCF-7 and MDA-MB-231 cells after mock treatment (0 nM Ipom-F) or treatment with 1 and 5 nM Ipom-F. Experiments were performed in at least three replicates using independently prepared biological samples. Panel (**B**) shows representative Western blot results for each protein. (**C**) Quantification of Western blot signal intensities for CD74, CST1, and CCN1. *p < 0.05, **p < 0.01, ***p < 0.001.

**Table 1.** Ipom-F-sensitive proteins identified in the time course and dose curve studies.

| UniProt ID | Protein Name | Gene ID | Number of Peptide | Number of Unique Peptide | Subcellular Localization | SILAC Ratio |  |  |  |
| --- | --- | --- | --- | --- | --- | --- | --- | --- | --- |
|  |  |  |  |  |  | Time course |  | Dose curve |  |
|  |  |  |  |  |  | 4h | 8h | 3nM | 6nM |
| Q16270 | Insulin-like growth factor-binding protein 7 | IGFBP7 | 7 | 7 | Secreted | 0.33 | 0.36 | 0.64 | 0.65 |
| P01037 | Cystatin-SN | CST1 | 6 | 3 | Secreted | 0.37 | 0.28 | 0.35 | 0.36 |
| P04233 | HLA class II histocompatibility antigen gamma chain | CD74 | 6 | 6 | membrane/<br>secreted | 0.42 | 0.24 | 0.46 | 0.35 |
| P01036 | Cystatin-S | CST4 | 6 | 2 | Secreted | 0.45 | 0.31 | 0.62 | 0.41 |
| O00622 | CCN family member 1 | CCN1 | 6 | 6 | Secreted | 0.54 | 0.41 | 0.49 | 0.34 |
| O43493 | Trans-Golgi network integral membrane protein 2 | TGOLN2 | 4 | 4 | Membrane | 0.61 | 0.43 | 0.60 | 0.40 |
| O95994 | Anterior gradient protein 2 homolog | AGR2 | 6 | 6 | ER/Secreted | 0.65 | 0.73 | 0.65 | 0.62 |
| O75976 | Carboxypeptidase D | CPD | 12 | 12 | Cell membrane | 0.70 | 0.41 | 0.56 | 0.53 |
| Q99519 | Sialidase-1 | NEU1 | 3 | 3 | Lysosome<br>membrane | 0.70 | 0.68 | 0.72 | 0.70 |
| P54709 | Sodium/potassium-transporting ATPase subunit beta-3 | ATP1B3 | 7 | 7 | Apical/Basolateral<br>membrane | 0.72 | 0.72 | 0.65 | 0.71 |
| P08648 | Integrin alpha-5 | ITGA5 | 7 | 7 | Cell membrane | 0.72 | 0.54 | 0.57 | 0.57 |
| Q03405 | Urokinase plasminogen activator surface receptor | PLAUR | 6 | 6 | Cell membrane/<br>secreted | 0.73 | 0.54 | 0.41 | 0.50 |

We selected three of the identified Ipom-F-sensitive proteins for expression verification. We treated triple-negative MDA-MB-231 cells with 0, 1, and 5 nM Ipom-F for 12 hours, and determined the expression of CD74 (a membrane/secreted protein), CST1 (a secreted protein), and CCN1 (a secreted protein) in the mock-treated and Ipom-F-treated cells using Western blotting. Consistent with the proteomic data (Table 1 and Supplementary Tables 3 and 4), Western blot analysis demonstrated that the levels of CD74, CST1, and CCN1 in MDA-MB-231 cells were significantly inhibited by 5 nM Ipom-F (Fig. 2B; compare lane 6 to lane 4; Fig. 2C). These results confirm that the method we used to select the Ipom-F-sensitive proteins was effective.

### Some of the identified Imp-F-sensitive proteins are upregulated in TNBC cells

To elucidate the expression of the 12 Ipom-F-sensitive proteins (Table 1) in various cancer cell lines, we utilized proteomics data from the Depmap portal (https://depmap.org/portal/) for each of these proteins in the NCI-60 human tumor cell lines (31). The portal provided comprehensive proteomics data on the expression of the 12 Ipom-F-sensitive proteins in 36 NCI-60 cancer panel cell lines. Our clustering analysis of the expression of these 12 proteins unveiled several major clusters among the 36 cell lines. Notably, one major cluster of cell lines expressed higher levels of PLAUR, ITGA5, CCN1, TGOLN2, CD74 and CST1 (Figure 3, top right corner). The second cluster showed lower levels of these proteins (Figure 3, top left corner). Intriguingly, the mesenchymal-like triple-negative MDA-MB-231 was in the first cluster, whereas two ERα-positive breast cancer cell lines, MCF-7 and T47D, were in the second cluster. These results suggest that the expression of PLAUR, ITGA5, CCN1, TGOLN2, CD74 and CST1 may play a role in differentiating the mesenchymal-like triple-negative MDA-MB-231 cells from other subtypes of cancer cells such as ERα-positive breast cancer cells. The expression of the remaining part of the 12 Ipom-F sensitive proteins, including NEU1, CPD, CST4, ATP1B3, IGFBP7, and AGR2 in the 36 cancer cell lines, was more sporadic. However, one notable feature was the elevated levels of most Ipom-F-sensitive proteins in MDA-MB-231 cells compared to the others (Fig. 3). Specifically, when examining the expression of the Ipom-F-sensitive proteins among the 36 NCI cancer cell lines, it was evident that the expression of 9 out of the 12 proteins (with CST1 lacking proteomic data in MDA-MB-231) was elevated in the MDA-MB-231 cells, making it the highest among the 36 cancer cell lines. Although proteomic data for CST1 in MDA-MB-231 cells was unavailable on the DepMap portal, our Western blot analysis demonstrated that CST1 protein levels in MDA-MB-231 cells were substantially higher than in MCF7 cells, similar to CD74 and CCN1 (Fig. 2B; compare lane 4 with lane 1).

**Figure 3.**
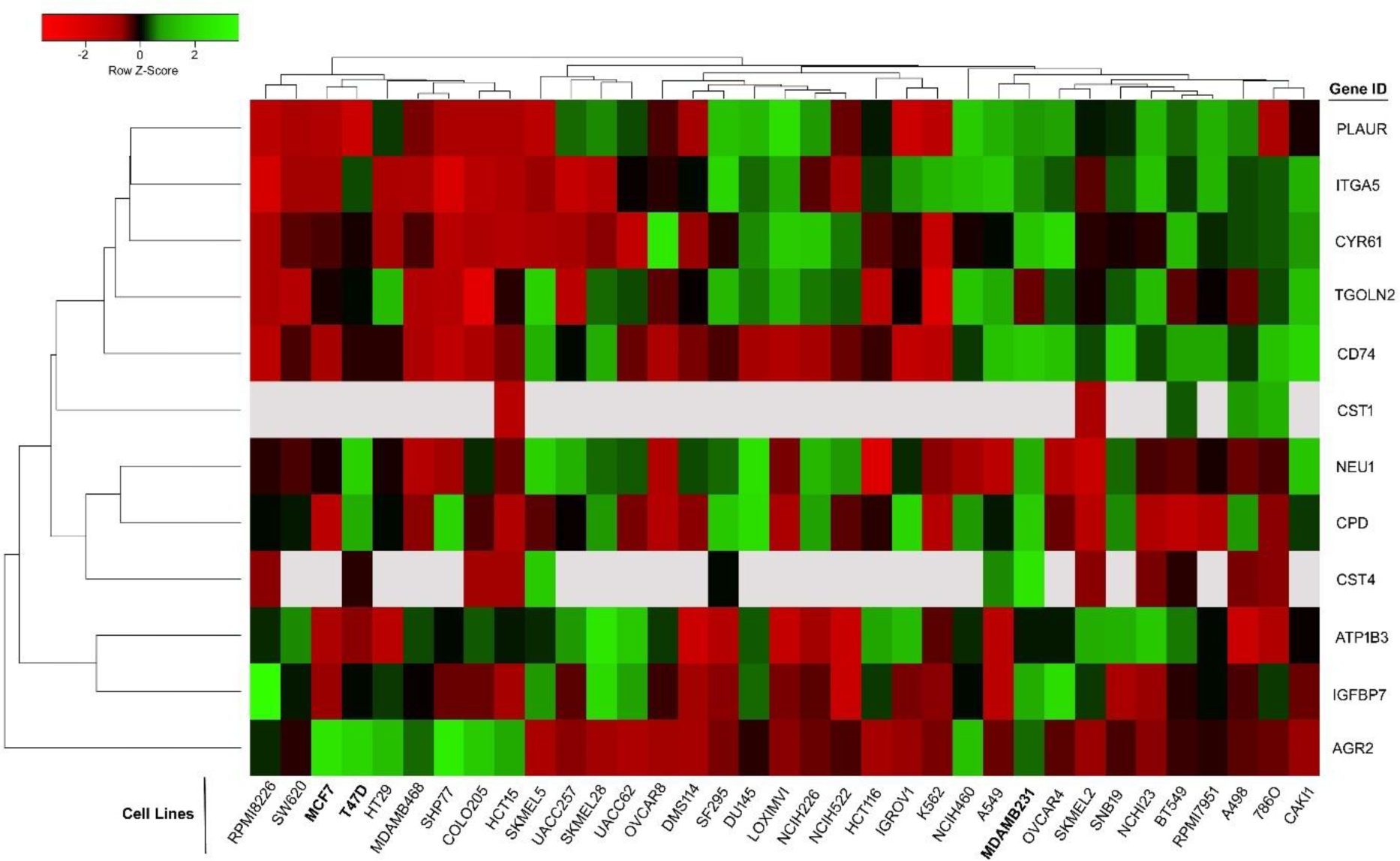
Expression levels of Ipom-F-sensitive proteins across 36 cells lines from the NCI-60 panel. Red = Downregulation; Green = Upregulation.

We also examined the protein levels of the Ipom-F-sensitive proteins in specimens from cancer patients using the UALCAN website (33, 34). The website contained proteomics data on some Ipom-F sensitive proteins, including CD74, PLAUR, ATP1B3, and CCN1, in various cancer specimens and normal tissues. CD74, PLAUR, and ATP1B3 were significantly elevated in TNBC tissues compared to luminal breast cancer and normal tissues (Fig. 4A-C). The CCN1 protein levels were significantly elevated in TNBC compared to normal tissues but not compared to luminal breast cancer, although the p-value was close to the threshold value for statistical significance Fig. 4D). We have verified the elevated levels of CD74 and the CCN1 protein in MDA-MB-231 cells compared to MCF-7 cells using Western blotting (Fig. 2B; compare lane 4 with lane 1). In summary, the results in this section suggest that the Ipom-F-sensitive proteins identified in this study are among the proteins upregulated in TNBC cells relative to normal cells and other subtypes of breast cancer cells. Based on these results, it is likely that the TNBC MDA-MB-231 cells are sensitive to Ipom-F because the cells have adapted to the elevated levels of these proteins, and once the expression of one or several of these proteins is inhibited by Ipom-F, the suppressed expression may lead to cell death and/or inhibition of proliferation.

**Figure 4.**
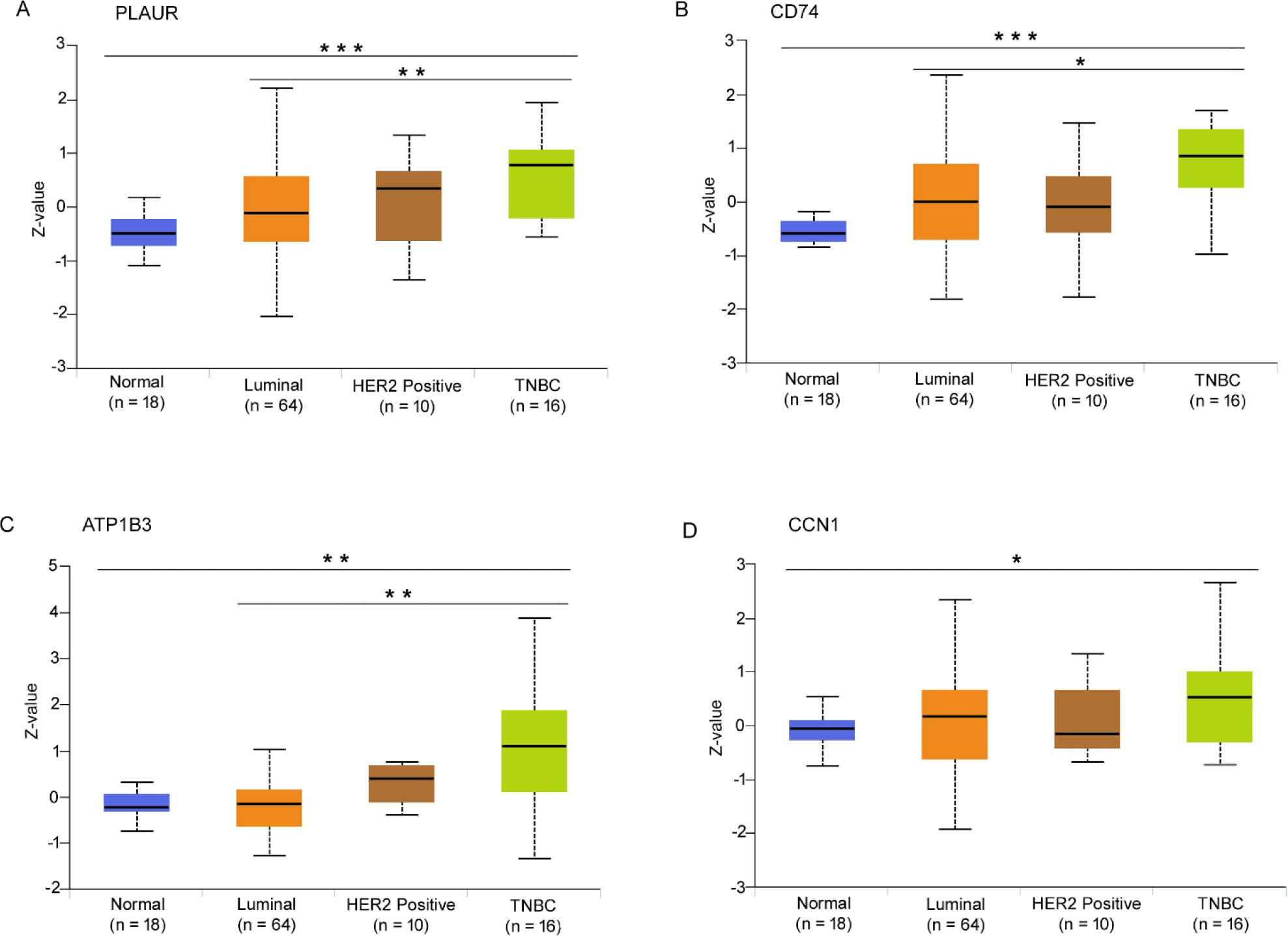
The expression levels of several Ipom-F-sensitive proteins in breast cancer tumor tissues. The protein levels of PLAUR (**A**) CD74 (**B**), (C), ATPIB3 (**C**), and CCN1 (**D**) in Luminal (n = 64), HER2-positive (n = 10), TNBC (n = 16), and normal breast tissue (n = 18). *\*p* ≤ *0.05; **p-value* ≤ *0.01; ***p-value* ≤ *0.001; n =* number of samples.

Although the import of proteins into the ER lumen and their subsequent transport to other cellular locations are conserved in all eukaryotic cells, Ipom-F appears to preferentially affect TNBC cells more severely than other cancerous and normal cells (7, 13). Drugs that disrupt conserved mechanisms in eukaryotic cells have been successfully used as chemotherapies (36, 37). For example, microtubule-targeting agents—chemical compounds that bind tubulin and interfere with microtubule assembly (e.g., colchicine) or disassembly (e.g., paclitaxel)—have been successfully used to treat various cancers (37, 38). Thus, Ipom-F has the potential to be used as a chemotherapy against TNBCs or other cancers, provided we gain a better understanding of the mechanisms underlying its cytotoxicity. This study represents one of the initial efforts toward that goal.

### Ipom-F potentially suppresses immune responses

To identify the functional categories of proteins affected by Ipom-F in the cells, we analyzed the Ipom-F-inhibited proteins across different time points and increasing doses of Ipom-F (Supplementary Tables 3 and 4) using FunRich (27, 28) and DAVID (29, 30). Interestingly, the functional enrichment analyses of the identified proteins indicated that Ipom-F significantly inhibited MHC class I and MHC class II receptor activities in both the time course (Fig. 5A) and dose curve (Fig. 5B) studies. The biological processes and molecular function enrichment analyses with DAVID confirmed the results obtained from the FunRich analysis (Supplementary Tables 5 and 6). MHC class I and II receptors play key roles in antigen presentation in the immune system, and the repression of MHC class I molecules leads to immune evasion by cancer cells (39, 40). Thus, while Ipom-F is being explored as an anti-cancer natural compound due to its cytotoxicity towards certain cancers (7, 13), its immunosuppressive effects, which can lead to immune evasion, cannot be completely ignored in cancer treatment exploration. On the other hand, the ability of Ipom-F to inhibit the expression of MHC class I and II molecules suggests that it may function as an immunosuppressant. Therefore, further investigations in this regard are warranted.

**Figure 5.**
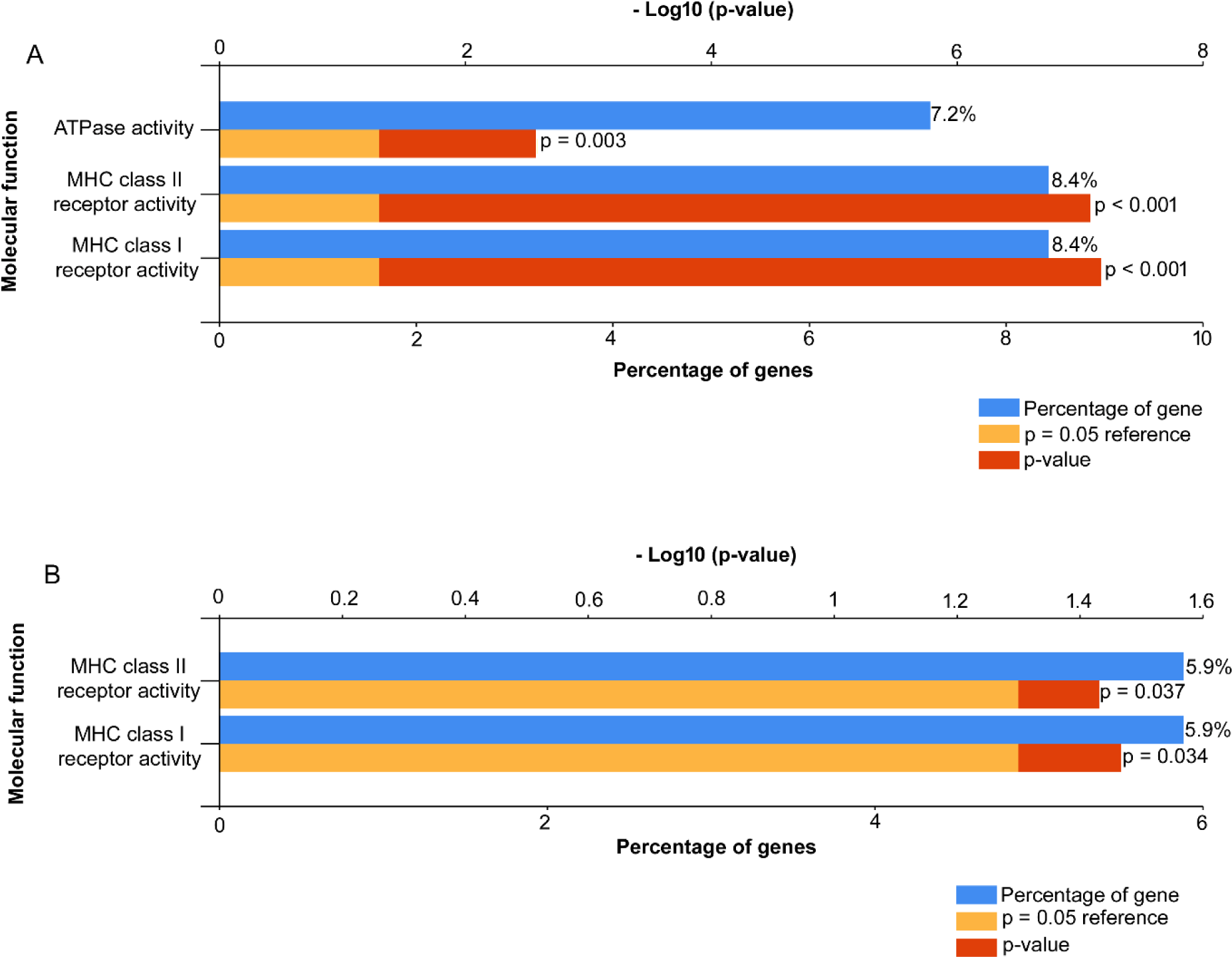
Ipom-F inhibits the protein levels of MHC class I and II molecules. The proteins that were consistently downregulated by Ipom-F across different time points (**A**) and increasing doses (**B**) were analyzed for enrichment of molecular functions using FunRich.

## Abbreviations

Ipom-F: ipomoeassin
F TNBC: triple-negative breast cancer

## Acknowledgements

This work was supported by Grant Number 2R15GM116032-02A1 from the National Institutes of Health (NIH) and a grant from Arkansas Biosciences Institute.

